# Antiviral activity of lambda-carrageenan against influenza viruses in mice and severe acute respiratory syndrome coronavirus 2 *in vitro*

**DOI:** 10.1101/2020.08.23.255364

**Authors:** Yejin Jang, Heegwon Shin, Myoung Kyu Lee, Oh Seung Kwon, Jin Soo Shin, Yongil Kim, Meehyein Kim

**Author notes:** **Corresponding author** Meehyein Kim, Ph.D., AdInfectious Diseases Therapeutic Research Center, Korea Research Institute of Chemical Technology, 141 Gajeongro, Yuseong, Daejeon 34114, Republic of Korea. siRNAgen Therapeutics Co., Daejeon 34302, Republic of Korea (Oh Seung Kwon).

## Abstract

Influenza virus and coronavirus, belonging to enveloped RNA viruses, are major causes of human respiratory diseases. The aim of this study was to investigate the broad spectrum antiviral activity of a naturally existing sulfated polysaccharide, lambda-carrageenan (λ-CGN), purified from marine red algae. Cell culture-based assays revealed that the macromolecule efficiently inhibited both influenza A and B viruses, as well as currently circulating severe acute respiratory syndrome coronavirus 2 (SARS-CoV-2), with EC_50_ values ranging from 0.3–1.4 μg/ml. No toxicity to host cells was observed at concentrations up to 300 μg/ml. Plaque titration and western blot analysis verified that λ-CGN reduced expression of viral proteins in cell lysates and suppressed progeny virus production in culture supernatants in a dose-dependent manner. This polyanionic compound exerts antiviral activity by targeting viral attachment to cell surface receptors and preventing entry. Moreover, intranasal administration to mice during influenza A viral challenge not only alleviated infection-mediated reductions in body weight but also protected 60% of mice from virus-induced mortality. Thus, λ-CGN could be a promising antiviral agent for preventing infection by several respiratory viruses.

## Introduction

Carrageenans (CGNs) extracted from marine seaweeds belong to a family of sulfated D-series polysaccharides harboring α-galactose residues. The diverse chemical structure and the degree of sulfation divides CGNs into three major polysaccharide groups, kappa (κ)-, iota (ι)- and lambda (λ)-CGNs, which contain one, two, and three negatively-charged sulfate ester groups per disaccharide repeating unit, respectively^1^. These natural polymers of diverse molecular weight have been used widely as pharmaceutical delivery vehicles that facilitate drug formulation or sustained drug release. As biomolecules, CGNs have various biological activities, including anticoagulant, anti-tumoral, or immunomodulatory functions ^2,3^. Several reports suggest that CGNs show *in vitro* or *in vivo* activity against rhinovirus, enterovirus 71, dengue virus, human herpes simplex, African swine fever virus, and influenza A virus ^4–10^. Most of these antiviral efficacy studies have focused on κ- and ι-CGNs; only one study suggested that λ-CGN was a potent inhibitor of rabies virus infection ^11^. Based on the structural characteristics of λ-CGN, by which it has no 3,6-anhydro-d-galactopyranosyl linkage as well as a higher sulfate content than the two other sulfated polysaccharides (Fig. 1A), we wondered whether it is active against two different respiratory viruses: influenza A and B viruses and severe respiratory syndrome coronavirus 2 (SARS-CoV-2).

**Figure 1.**
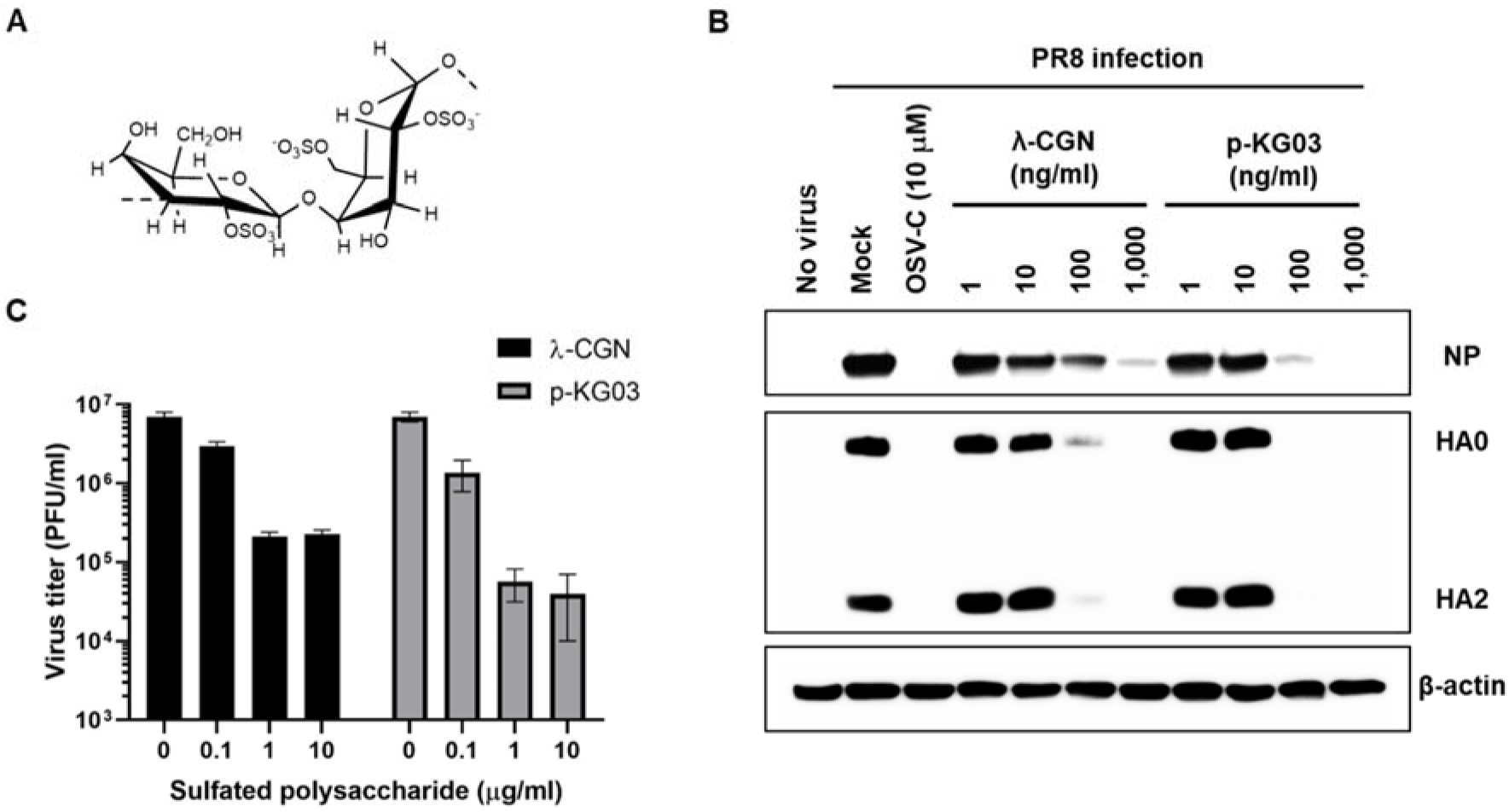
λ-CGN inhibits influenza virus infection *in vitro*. (A) Chemical structure of the repeating disaccharide unit in λ-CGN. (B) Western blot analysis showing expression of viral proteins. MDCK cells infected with PR8 at an MOI of 0.001 were mock-treated (Mock) or treated with increasing concentrations of λ-CGN or p-KG03, or with 10 μM of OSV-C, at 35°C. On the next day, cell lysates were harvested for SDS-PAGE and immunoblotting with anti-NP or anti-HA antibodies. β-Actin was used as a loading control. ‘No virus’ means negative control without viral infection. (C) Plaque assay to determine viral titers. Ten-fold serial dilutions of cell culture supernatants acquired after infection and compound treatment in (B) were loaded onto fresh MDCK cells and cultured at 33°C in 1.2% Avicel-containing overlay medium. The number of viral plaques was counted after crystal violet staining on day 3 p.i. Data are expressed as the mean ± SEM of three independent experiments.

Influenza virus is a major human respiratory virus that causes seasonal epidemics or unexpected pandemic outbreaks. It belongs to the family *Orthomyxoviridae* and contains an eight-segmented, negative-sense RNA genome classified into three types, A, B and C. Type A is further divided into subtypes based on the serological characteristics of surface glycoproteins hemagglutinin (HA) and neuraminidase (NA), while type B is split into Victoria and Yamagata lineages. Even though therapeutic antivirals such as oseltamivir phosphate, zanamivir, peramivir and baloxavir marboxyl, as well as preventative vaccines, have been successfully developed, emerging drug-resistant strains and mismatch-derived inefficacy of vaccines mean that this virus remains a threat to human public health, with an estimated annual mortality burden of 290,000 to 650,000 deaths ^12–14^.

Coronavirus, a member of the family *Coronaviridae*, is also an enveloped virus with a positive-sense single-stranded RNA genome of 26 to 32 kilobases in length. Similar to influenza virus, it is a zoonotic virus that causes respiratory disease in humans. Most infections cause mild symptoms such as fever, fatigue, or dry cough. However, recently emerging viruses have become more lethal and highly contagious. For example, SARS-CoV, first identified in 2003, had a mortality rate of 10%, with over 8,000 laboratory-confirmed cases, whereas Middle East respiratory syndrome coronavirus (MERS-CoV), identified in 2012, had a mortality rate of 34%, with 2,494 cases ^15,16^. In comparison to SARS-CoV and MERS, currently circulating SARS-CoV-2 has a lower fatality rate (about 9% compared with SARS-CoV-1); however, it has caused a global pandemic, with over fourteen million confirmed cases and 607 thousand deaths recorded since December 2019 ^17^. Despite this formidable circulation, we still have no coronavirus-specific antivirals or vaccines. Because the symptoms and transmission routes of these respiratory viruses are very similar, a broad-spectrum antiviral agent is required for their co-treatment. Therefore, the aim of this study was to assess the antiviral activity of λ-CGN against influenza viruses and SARS-CoV-2 and to identify the mechanism of action.

## Experimental Section

### Cells, viruses, and compounds

Madin-Darby canine kidney (MDCK) and African green monkey kidney cells (Vero) were purchased from the American Type Culture Collection (Cat. Nos., CCL-34 and CCL-81; ATCC, Manassas, VA, USA). They were maintained in minimum essential medium (MEM; HyClone, Logan, UT, USA) and Dulbecco’s modified Eagle’s medium (DMEM; HyClone), respectively, supplemented with 10% fetal bovine serum (FBS; Atlas Biologicals, Fort Collins, CO, USA). Influenza viruses A/Puerto Rico/8/34 (PR8; H1N1), A/Hong Kong/8/68 (HK; H3N2), and B/Lee40 (Lee) were purchased from the ATCC. The mouse-adapted PR8 (maPR8) strain was a kind gift from Prof. H. J. Kim (Chung-Ang University, Seoul, Republic of Korea). Influenza A viruses were inoculated into 10-day-old embryonated chicken eggs at 37°C for 3 days, whereas influenza B virus was amplified at 35°C for 3 days in MDCK cells in the presence of 2 μg/ml tosyl phenylalanyl chloromethyl ketone (TPCK)-treated trypsin (Sigma-Aldrich, St. Louis, MO, USA). SARS-CoV-2 (BetaCoV/Korea/KCDC03/2020), provided by Korea Centers for Disease Control and Prevention, was amplified in Vero cells at 37°C for 3 days. After centrifugation at 1,000 *g* for 5 min, viral stocks were stored at −80°C and viral titers were determined in a plaque assay using crystal violet ^18^. The test compound λ-CGN, average molecular weight 1,025 kDa, was purchased from DuPont Nutrition & Biosciences (Wilmington, DE, USA). Control anti-influenza viral agents amantadine hydrochloride (AMT; ≥98%) and ribavirin (RBV; ≥98%) were purchased from Sigma-Aldrich. Oseltamivir carboxylate (OSV-C) was purchased from United States Biological (Swampscott, MA, USA). Marine microalgae-derived sulfated polysaccharide p-KG03 was provided and characterized by Dr. Joung Han Yim (Korea Polar Research Institute, Incheon, Republic of Korea) ^19^. Oseltamivir phosphate (OSV-P; ≥98%) for *in vivo* antiviral studies was obtained from Hanmi Pharmaceutical Co. (Gyeonggi-do, Republic of Korea). Remdesivir (RDV; 99.74%), a control anti-SARS-CoV-2 compound, was purchased from MedChem Express (Monmouth Junction, NJ, USA).

### Cell culture-based antiviral assay

An antiviral assay for influenza viruses was performed as described previously ^20^. Briefly, MDCK cells grown overnight in 96-well plates (3 × 10^4^ cells per well) were mock-infected or infected with each viral strain at a multiplicity of infection (MOI) of 0.001 at 35°C for 1 h. After removing unabsorbed virus, cells were treated with 3-fold dilutions of each compound for 3 days at the same temperature. Viability of non-infected or infected cells was measured using 3-(4,5-dimethylthiazol-2-yl)-2,5-diphenyltetrazoliumbromide (MTT) to determine the half-maximal cytotoxic concentration (CC_50_) and the half-maximal effective concentration (EC_50_), respectively. To assess anti-SARS-CoV-2 activity, Vero cells were grown overnight in 96-well plates (2 × 10^4^ cells per well). After addition of compounds (serially diluted 3-fold), cells were infected at 37°C for 2 days with an equal volume of SARS-CoV-2 (MOI of 0.05) in a biosafety level 3 laboratory. The cells were fixed and permeabilized with chilled acetone:methanol (1:3) for probing with an anti-S antibody (Genetex, Irvine, CA) followed by Alexa Fluor 488-congujated goat anti-mouse IgG (Invitrogen, Carlsbad, CA) to determine EC_50_ values. Cell nuclei were counterstained with 4′,6-diamidino-2-phenylindole (DAPI; Invitrogen) to calculate the CC_50_ values. The number of S-derived (green) and cell nuclei-derived (blue) signals detected in four spots per well from three independent samples was quantified using the Operetta high content screening system (Perkin Elmer, Waltham, MA, USA) and the built-in Harmony software.

### Western blot analysis

PR8-infected MDCK cells (MOI, 0.001) were treated with increasing concentrations of λ-CGN, pKG-03 or OSV-C at 35°C. On the next day, culture lysates were harvested and loaded onto 10 or 12% SDS-PAGE gels (40 μg total protein per well) for electrotransfer. Viral NP and HA proteins were detected using mouse anti-NP (catalog no. 11675-MM03; Sino Biological, Beijing, China) and rabbit anti-HA2 (catalog no. 86001-RM01; Sino Biological) antibodies, respectively, according to our previous report ^18^. Cellular β-actin was used as a loading control and detected using a mouse anti-β-actin antibody (catalog no. A1987; Sigma-Aldrich). Horseradish peroxidase (HRP)-conjugated goat anti-mouse or anti-rabbit secondary antibodies were used to detect the primary antibodies (Thermo Scientific, Waltham, MA, USA). After addition of a chemiluminescent HRP substrate (SuperSignal West Pico Chemiluminescent Substrate; Pierce, Rockford, IL, USA), images were obtained using a LAS-4000 Luminescent Image Analyzer (Fujifilm, Tokyo, Japan).

### Plaque titration

A plaque assay was performed as described previously, with some modifications ^18^. Briefly, MDCK cells seeded in 6-well plates were infected with PR8 at an MOI of 0.001 in the absence or presence of increasing concentrations of λ-CGN or p-KG03. On the next day, the culture supernatants were harvested and 10-fold serial dilutions were used to infect fresh MDCK cells. After incubation of infected cells in overlay medium [serum-free MEM with 1.2% Avicel RC-591 (FMC Corp, Philadelphia, PA, USA) and 2 μg/ml TPCK-trypsin (Sigma-Aldrich)] at 33°C for 3 days, the number of plaques was counted by crystal violet staining.

### Confocal microscopy

PR8-infected MDCK cells were mock-treated or treated with the sulfated polysaccharides (10 μg/ml) for 4 h at 37°C. In parallel, the same samples were incubated for 2.5 h at 37°C with protein synthesis inhibitor cycloheximide (10 μg/ml) (CHX; Sigma-Aldrich), of which experimental condition was optimized in our previous reports ^18,21^. Viral NP was visualized using an anti-NP antibody (cat no. sc-80481; Santa Cruz Biotechnology) and Alexa Fluor 488-conjugated goat anti-mouse IgG (Invitrogen), while nuclear DNA was counterstained using 4′,6-diamidino-2-phenylindole (DAPI; Vector Laboratories, Burlingame, CA, USA). Images were captured under a Zeiss LSM 700 confocal microscope and data were analyzed using ZEN software (Carl Zeiss, Thornwood, NY, USA).

### *In vivo* study

Antiviral efficacy study in a mouse model was performed by modification of our previous report ^18^. Brifly, female BALB/c mice (6–7 weeks old; Orient Bio Inc., Gyeonggi-do, Republic Korea) were infected with maPR8. Five units of 50% mouse lethal dose (5 MLD_50_) of the virus were preincubated with λ-CGN for 30 min at room temperature. Mice were challenged intranasally with maPR8 alone or with maPR8 mixed with λ-CGN (1 or 5 mg/kg) in a total volume of 50 μl. The control group received OSV-P from days 0 to 5 p.i. (10 mg/kg/day (b.i.d.)) beginning 4 h before virus challenge. Changes in body weight and mortality were measured every day for 15 days. Mice were sacrificed when they lost at least 25% of their body weight. All animal experiments were conducted in accordance with ethical guidelines approved by the Institutional Animal Care and Use Committee (IACUC) of the Korea Research Institute of Chemical Technology (KRICT). All experimental procotols were approved by the KRICT’s IACUC with the code number of 2020-6D-04-01. Kaplan–Meier survival curves were constructed using GraphPad Prism 6 (GraphPad Software, San Diego, CA).

## Results

### Anti-influenza activity of λ-CGN

To examine the antiviral activity of λ-CGN, increasing concentrations of the compound were used to treat influenza virus-infected MDCK cells. Another sulfated polysaccharide, p-KG03, of which antiviral activity has been elucidated in our previous report ^19^and three different antiviral chemicals (AMT, RBV and OSV-C) were used as controls. The anti-influenza viral activity or the drug-resistance profiles of all of these control compounds were reproducible, indicating that the cell culture-based antiviral assay is reliable. The CPE assay on day 3 p.i. revealed that λ-CGN efficiently inhibited infection by both influenza A and B viruses, with EC_50_ values of 0.3 to 1.4 μg/ml, with no cytotoxicity up to a maximum concentration of 300 μg/ml (Table 1). Notably, the inhibitory effect was comparable with that of p-KG03. To confirm this finding, we measured changes in viral protein expression in cell lysates and infectious viral titers in culture supernatants (Fig. 1B and C, Supplementary Fig. S1). The data revealed not only that λ-CGN is able to inhibit expression of viral proteins NP and HA in infected cells, but also that is suppressed production of progeny virus in a dose-dependent manner as in the p-KG03-treated samples. These results suggest that λ-CGN has potent antiviral activity against influenza A and B viruses *in vitro*, with selectivity index (SI) values over 214.3.

**Table 1.**
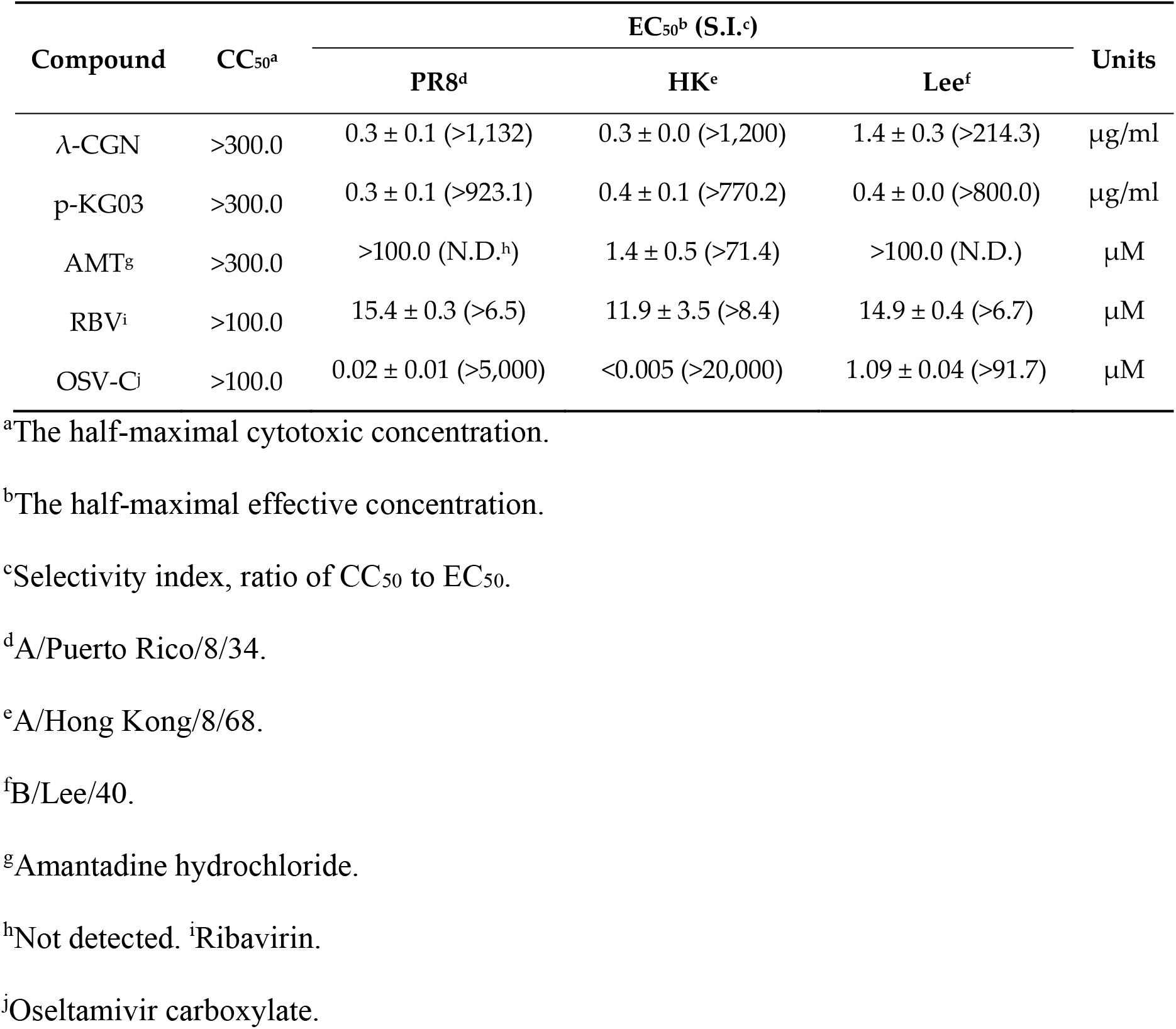
Antiviral effect of λ-CGN against influenza A and B viruses

### λ-CGN inhibits influenza viral entry

We wondered which step of the influenza virus life cycle is targeted by λ-CGN. Intracellular distribution of viral NP was first compared in the absence or presence of the compound at 4 h p.i, a time when NP was fully localized to the nuclei accompanied by robust replication of viral RNA (Fig. 2A). The confocal microscopic images revealed that, similar to p-KG03, λ-CGN reduced the number of NP-positive nuclei when compared to the mock compound-treated sample. However, inhibition by the two polymers had little effect on NP-derived fluorescent intensity. This finding during a single round of infection suggests that λ-CGN targets the virus entry step rather than RNA-dependent RNA replication or viral protein expression. To clarify its mode of action, we monitored the intracellular distribution of NP at an earlier time point (2.5 h p.i.,) in the presence of CHX, a protein synthesis inhibitor that allows tracking of the input viral protein and its localization. Under these conditions, when NP was present in the cytoplasm but not reaching the nucleus, λ-CGN completely blocked membrane penetration of the viral particles harboring vRNP complexes as efficiently as p-KG03. No NP accumulated on the surface of the cellular membrane strongly suggests that λ-CGN targets attachment of influenza virus to its cell surface receptors.

**Figure 2.**
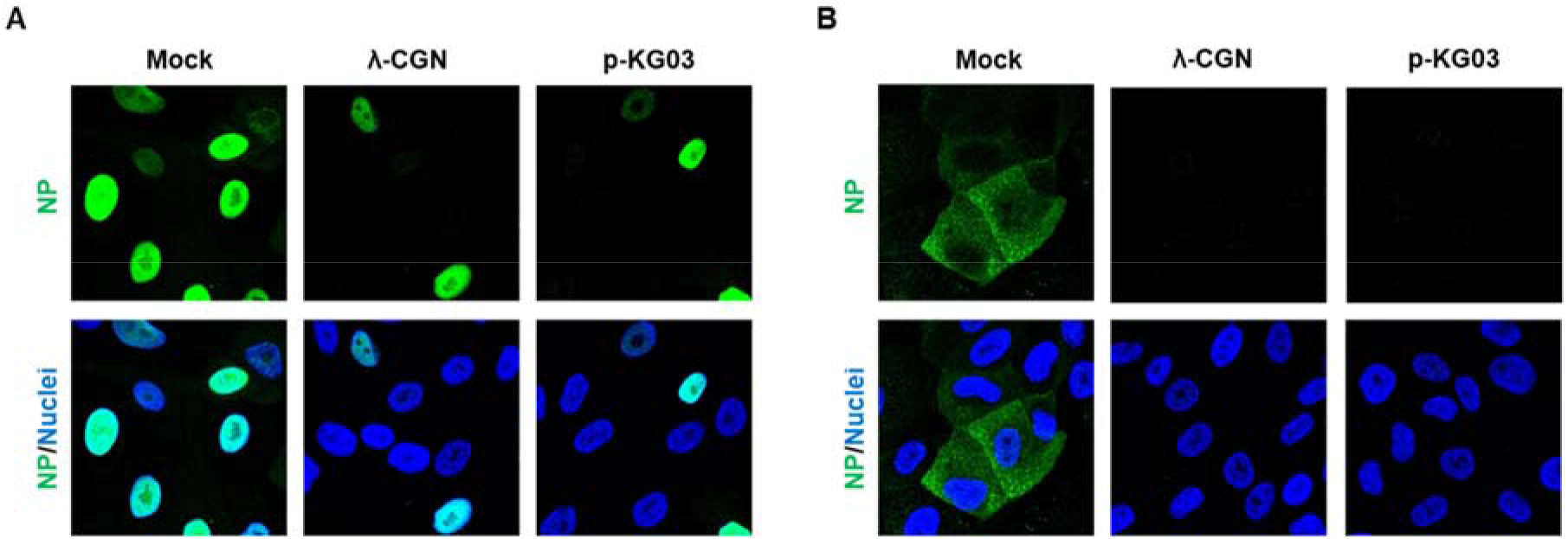
Effect of λ-CGN on the intracellular entry of influenza A virus. MDCK cells were infected with PR8 (MOI, 5) and subsequently mock-treated or treated either with λ-CGN or with p-KG03 at a concentration of 10 μg/ml. At 4 h p.i. in the absence of CHX (A) or at 2.5 h in the presence of 10 μg/ml CHX (B), viral NP was detected with an anti-NP antibody and an Alex Fluor 488-conjugated goat anti-mouse secondary antibody (green). Cell nuclei were counterstained with DAPI (blue). Original magnification, 400×.

### λ-CGN protects mice from infection by influenza virus

To investigate the antiviral activity of λ-CGN *in vivo*, mice were infected intranasally with maPR8 alone or with maPR8 plus λ-CGN once. As a control, maPR8-infected mice received oral OSV-P twice a day for 6 days. Antiviral activity was determined by monitoring body weight and mortality for 15 days. The results revealed that maPR8 at 5 MLD_50_ caused body weight loss (Fig. 3A) and complete death (Fig. 3B) within 8 days. Interestingly, intranasal administration of 5 mg/kg λ-CGN abrogated infection-mediated body weight loss, yielding a 60% survival rate for infected mice. However, this antiviral efficacy was not observed at a lower dose (1 mg/kg). As expected, treatment with OSV-P at 10 mg/kg/day for 6 days showed notable therapeutic effects, ensuring the reliability of the *in vivo* antiviral study. Taken together, these data suggest that intranasal co-administration of λ-CGN prevents viral infection-mediated body weight loss and reduces mortality.

**Figure 3.**
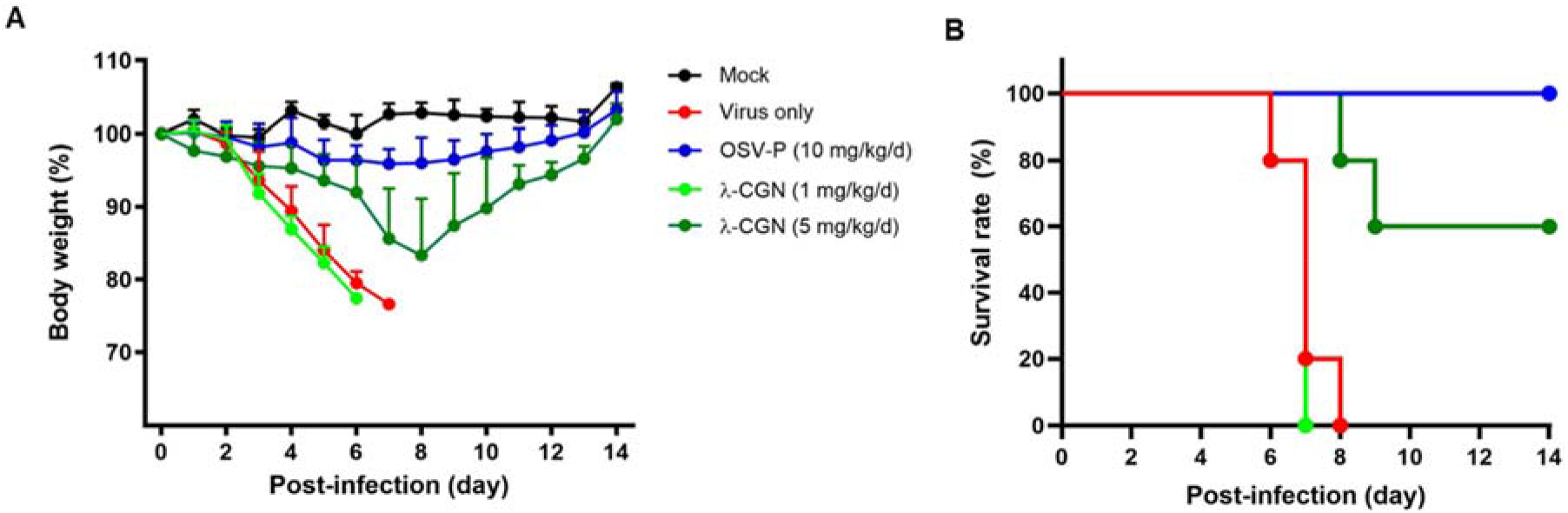
Effect of λ-CGN on influenza A infection *in vivo*. BALB/c mice were mock-infected (black) or intranasally infected with maPR8 at 5 MLD50 (red). As test groups, the virus was preincubated at room temperature for 30 min with λ-CGN at a lower dose (1 mg/kg/d, bright green) or a higher dose (5 mg/kg/d, dark green), followed by intranasal administration. Control mice received oral OSV-P twice a day (10 mg/kg/d) at 8-h intervals, starting at 4 h before viral infection (blue). Body weight (A) and mortality (B) of mice were measured every day from Days 0 to 14 post-infection. Data are expressed as the mean ± SEM from five mice

### Anti-SARS-CoV-2 activity of λ-CGN

Next, we asked whether the sulfated polysaccharide has antiviral activity against another enveloped respiratory virus, SARS-CoV-2. Vero cells infected with the virus at an MOI of 0.1 were treated with increasing concentrations of λ-CGN by using RDV as a control. On Day 2, immunofluorescence staining with an anti-viral S antibody revealed that SARS-CoV-2 infection was inhibited effectively by the test compound, without affecting cell viability (Figure 4A). As expected, anti-SARS-CoV-2 activity was well visualized in the RDV-treated cells. Quantitative analysis of antiviral dose-response and cell viability showed that λ-CGN had an EC_50_ of 0.9 ± 1.1 μg/ml and a CC_50_ of >300.0 μg/ml (resulting in an S.I., > 333.3), while RDV had an EC_50_ of 23.5 ± 1.2 μM and a CC_50_ of >300.0 μM (resulting in an S.I., >12.8). These results demonstrate that λ-CGN is highly active against SARS-CoV-2.

**Figure 4.**
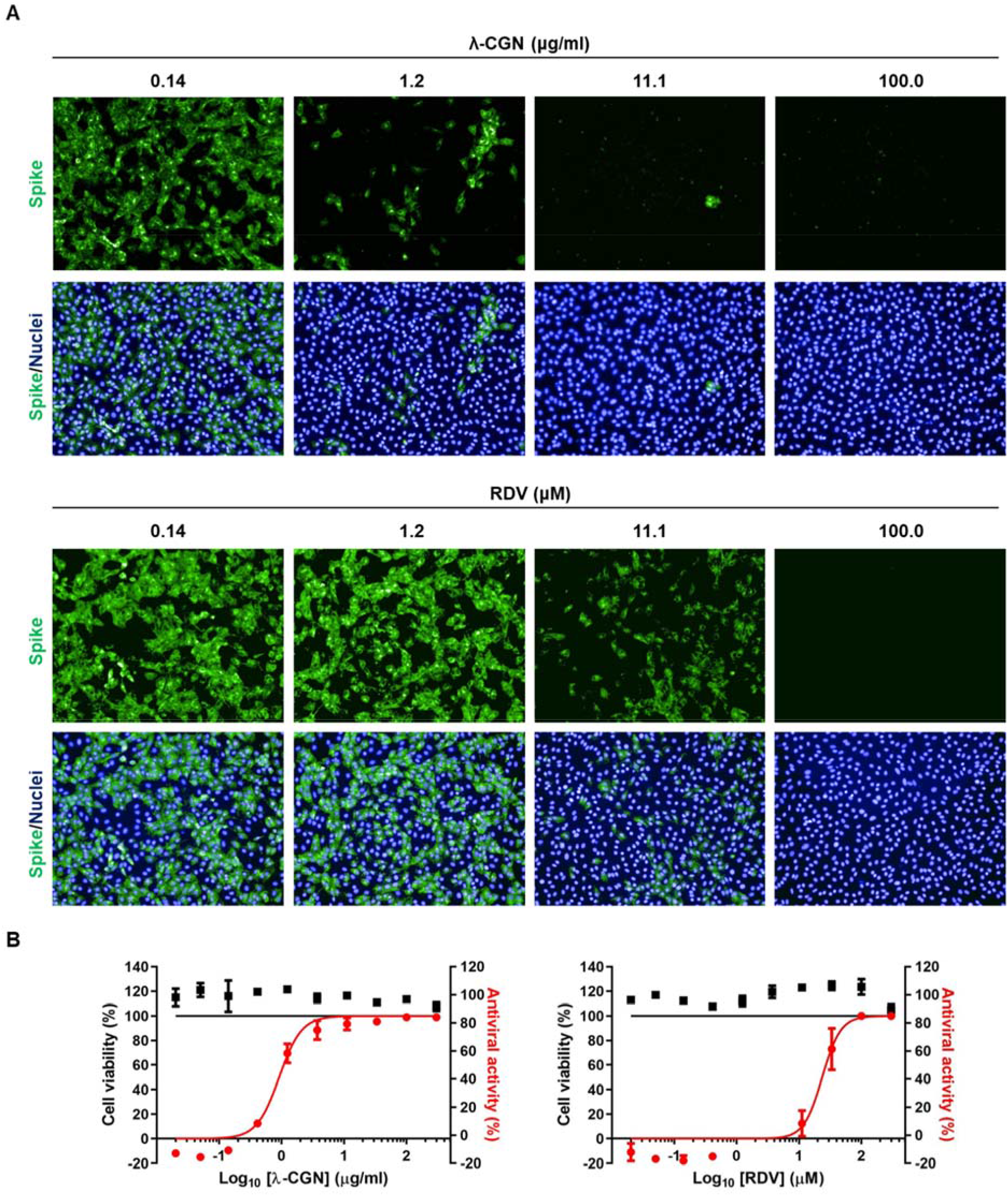
Anti-SARS-CoV-2 activity of λ-CGN. (A) Vero cells seeded in 96-well plates were infected with SARS-CoV-2 at an MOI of 0.1, either alone or in the presence of increasing concentrations of λ-CGN (upper panel) or RDV (lower panel; a control). On Day 2 post-infection, cells were fixed and permeabilized prior to immunostaining with an anti-viral S antibody and an Alexa Fluor 488-conjugated goat anti-mouse IgG (green). Cell nuclei were counterstained with DAPI to estimate cell viability (blue). Images were captured with a 20× objective lens fitted to an automated fluorescence microscope (Operetta HCS). (B) The number of fluorescent spots was counted to calculate antiviral activity (green) and cell viability (blue) at each concentration of the compounds. The viability of mock-infected cells were fixed as 100%, while the antiviral activity in virus-infected cells or mock-infected cells was fixed as 0 and 100%, respectively. Data are expressed as the mean ± SEM from independent experiments.

## Discussion

Sulfated polysaccharides such as heparin, dextran sulfate, and pentosan sulfate, as well as various CGNs, show antiviral or virucidal activity against diverse enveloped viruses at subtoxic concentrations ^22–26^. These studies of the physiochemical properties and molecular structure of these compounds reveal that their antiviral efficacy is mainly due to their affinity for viral glycoproteins, resulting in blockade of viral attachment to cellular receptors; the charge density, chain length, degree of sulfation, and detailed structural features of these macromolecules are critical for this interaction. In-depth studies of the underlying mechanisms demonstrate that the macromolecules exert anti-HIV activity by competing with polyanionic regions of host-cell-surface molecules for binding to the positively charged amino acids present in the viral enveloped glycoprotein, gp120, including the V3 loop ^27–29^. The microbicidal activity of polystyrene sulfonate against sexually transmitted infectious diseases caused by HSV-2 and papillomavirus has been evaluated *in vivo* and *in vitro* ^30,31^. Unfortunately, prevention of vaginal HIV transmission using topical cellulose sulfate gel failed ^32^, indicating the need to develop a more potent microbicidal sulfated polysaccharide or to administer other polymers through an alternative route, such as oral or intranasal.

Regarding this issue, it is not strange to anticipate that intranasal treatment with active sulfated polysaccharides could be a promising way to prevent infection by various respiratory enveloped viruses such as influenza A and B viruses, respiratory syncytial virus, and coronaviruses. Previously, it was reported that κ-CGN with a molecular weight of 2 kDa is active against influenza A virus *in vitro*, with an EC_50_ value of 32.1 μg/ml. In addition, ι-CGN inhibited influenza A virus infection of MDCK cells with an EC_50_ value of 0.04–0.20 μg/ml; not only that, intranasal administration of ι-CGN showed therapeutic effects in an influenza A virus-infected mouse model ^4,33^. Notably, a randomized double-blind study in volunteers with early symptoms of the common cold confirmed the efficacy and safety of an antiviral ι-CGN nasal spray ^34^. In contrast to κ- and ι-CGNs, the antiviral activity of λ-CGN has rarely been investigated in the context of viruses that are transmitted in droplets or through the air. Therefore, we wondered whether λ-CGN is able to inhibit both influenza A and B viruses and/or the emerging coronavirus SARS-CoV-2. We were interested in λ-CGN, because this compound comprises alternating (1,3)-linked α-D-galactose-2-sulated and (1,4)-linked β-D-galactose-2,6,-disulfated units, has a higher degree sulfation with an ester sulfate content of about 32– 39%, and shows better solubility in cold water than the other two CGNs ^35^. Accordingly, the sulfated polysaccharide was expected to have efficient and broad antiviral activity and to be easily dissolved in an aqueous solution when it is formulated for a nasal spray.

Similar to the other sulfated polysaccharides mentioned above, we observed that λ-CGN targets the influenza virus entry step. Strikingly, its virucidal properties led to a 60% survival rate in virus-challenged mice after an exposure of infectious virus to the antiviral agent. However, it is unclear whether this polyanionic compound is able to protect small animals such as hACE-expressing mice or Syrian hamsters from SARS-CoV-2 infection by blocking the viral S protein-associated entry step ^36,37^. In addition, because CGNs have intrinsic anti-coagulant activity, any unwarranted side effects should be ruled out before clinical application. This is because dysfunctional or aberrant coagulation is responsible for the hyper-inflammatory responses observed in severe cases of influenza or SARS-CoV-2 infection-mediated pneumonia, and anti-coagulant signals could be over-stimulated already in the lungs of infected patients ^38,39^. To the best of our knowledge, this is the first report to suggest that λ-CGN inhibits infection by influenza B as well as influenza A viruses and emerging SARS-CoV-2. The broad spectrum antiviral activity of this compound would make it effective against different families of respiratory virus that are circulating concurrently and when prophylactic treatment is definitely required before diagnosis.

## Acknowledgements

The authors thank Dr. Joung Han Yim at the Korea Polar Research Institute for providing p-KG03. The SARS-CoV-2 resource (NCCP No., 43326) for this study was provided by the National Culture Collection for Pathogens. This research was supported by the National Research Foundation of Korea (NRF) grants funded by the Korea government (MSIT) (grant number NRF-2020M3A9I2081687 to M.K.) and by an intramural fund from KRICT (grant number KK1703-E00 to M.K.).

## Author contributions

M.K. wrote and edited the main manuscript text. M.K. and Y.J. planned and supervised the experiments. Y.J., H.S., O.S.K. and J.S.S. performed the antiviral assays using infectious viruses. Y.J. and M.K.L. analyzed and visualized the data. Y.K. prepared and characterized the carrageenan. All authors reviewed the manuscript.

## Additional information

Y.J., H.S., M.K.L, O.S.K., J.S.S. and M.K. declare no conflict of interest. Y.K. is trying to commercialize λ-CGN used in this study through his company Hanmi Pharmaceutical Co.

## References

1 Liu, J., Zhan, X., Wan, J., Wang, Y. & Wang, C. Review for carrageenan-based pharmaceutical biomaterials: favourable physical features versus adverse biological effects. Carbohydr Polym 121, 27–36, doi:10.1016/j.carbpol.2014.11.063 (2015).

2 Yao, Z., Wu, H., Zhang, S. & Du, Y. Enzymatic preparation of kappa-carrageenan oligosaccharides and their anti-angiogenic activity. Carbohydr Polym 101, 359–367, doi:10.1016/j.carbpol.2013.09.055 (2014).

3 Liang, W., Mao, X., Peng, X. & Tang, S. Effects of sulfate group in red seaweed polysaccharides on anticoagulant activity and cytotoxicity. Carbohydr Polym 101, 776–785, doi:10.1016/j.carbpol.2013.10.010 (2014).

4 Wang, W. et al. In vitro inhibitory effect of carrageenan oligosaccharide on influenza A H1N1 virus. Antiviral Res 92, 237–246, doi:10.1016/j.antiviral.2011.08.010 (2011).

5 Gonzalez, M. E., Alarcon, B. & Carrasco, L. Polysaccharides as antiviral agents: antiviral activity of carrageenan. Antimicrob Agents Chemother 31, 1388–1393, doi:10.1128/aac.31.9.1388 (1987).

6 Talarico, L. B. et al. The antiviral activity of sulfated polysaccharides against dengue virus is dependent on virus serotype and host cell. Antiviral Res 66, 103–110, doi:10.1016/j.antiviral.2005.02.001 (2005).

7 Talarico, L. B., Noseda, M. D., Ducatti, D. R. B., Duarte, M. E. R. & Damonte, E. B. Differential inhibition of dengue virus infection in mammalian and mosquito cells by iota-carrageenan. J Gen Virol 92, 1332–1342, doi:10.1099/vir.0.028522-0 (2011).

8 Chiu, Y. H., Chan, Y. L., Tsai, L. W., Li, T. L. & Wu, C. J. Prevention of human enterovirus 71 infection by kappa carrageenan. Antiviral Res 95, 128–134, doi:10.1016/j.antiviral.2012.05.009 (2012).

9 Grassauer, A. et al. Iota-Carrageenan is a potent inhibitor of rhinovirus infection. Virol J 5, 107, doi:10.1186/1743-422X-5-107 (2008).

10 Garcia-Villalon, D. & Gil-Fernandez, C. Antiviral activity of sulfated polysaccharides against African swine fever virus. Antiviral Res 15, 139–148, doi:10.1016/0166-3542(91)90031-l (1991).

11 Luo, Z. et al. lambda-Carrageenan P32 Is a Potent Inhibitor of Rabies Virus Infection. PLoS One 10, e0140586, doi:10.1371/journal.pone.0140586 (2015).

12 Hayden, F. G. & de Jong, M. D. Emerging influenza antiviral resistance threats. J Infect Dis 203, 6–10, doi:10.1093/infdis/jiq012 (2011).

13 Andres, C. et al. Molecular influenza surveillance at a tertiary university hospital during four consecutive seasons (2012-2016) in Catalonia, Spain. Vaccine 37, 2470–2476, doi:10.1016/j.vaccine.2019.03.046 (2019).

14 Paget, J. et al. Global mortality associated with seasonal influenza epidemics: New burden estimates and predictors from the GLaMOR Project. J Glob Health 9, 020421, doi:10.7189/jogh.09.020421 (2019).

15 Centers for Disease Control and Prevention. In the Absence of SARS-CoV Transmission Worldwide: Guidance for Surveillance, Clinical and Laboratory Evaluation, and Reporting. https://www.cdc.gov/sars/surveillance/absence.html (2005).

16 World Health Organization. Middle East respiratory syndrome coronavirus (MERS-CoV). https://www.who.int/emergencies/mers-cov (2019).

17 World Health Organization. Coronavirus disease 2019 (COVID-19) Situation Report–193. https://www.who.int/docs/default-source/wha-70-and-phe/20200721-covid-19-sitrep-183.pdf?sfvrsn=b3869b3_2 (2020).

18 Jang, Y. et al. Salinomycin Inhibits Influenza Virus Infection by Disrupting Endosomal Acidification and Viral Matrix Protein 2 Function. J Virol 92, doi:10.1128/JVI.01441-18 (2018).

19 Kim, M. et al. In vitro inhibition of influenza A virus infection by marine microalga-derived sulfated polysaccharide p-KG03. Antiviral Res 93, 253–259, doi:10.1016/j.antiviral.2011.12.006 (2012).

20 Jang, Y. et al. In Vitro and In Vivo Antiviral Activity of Nylidrin by Targeting the Hemagglutinin 2-Mediated Membrane Fusion of Influenza A Virus. Viruses 12, doi:10.3390/v12050581 (2020).

21 Jang, Y. et al. Antiviral activity of KR-23502 targeting nuclear export of influenza B virus ribonucleoproteins. Antiviral Res 134, 77–88, doi:10.1016/j.antiviral.2016.07.024 (2016).

22 Nyberg, K. et al. The low molecular weight heparan sulfate-mimetic, PI-88, inhibits cell-to-cell spread of herpes simplex virus. Antiviral Res 63, 15–24, doi:10.1016/j.antiviral.2004.01.001 (2004).

23 Mitsuya, H. et al. Dextran sulfate suppression of viruses in the HIV family: inhibition of virion binding to CD4+ cells. Science 240, 646–649, doi:10.1126/science.2452480 (1988).

24 Andrei, G., Snoeck, R., Goubau, P., Desmyter, J. & De Clercq, E. Comparative activity of various compounds against clinical strains of herpes simplex virus. Eur J Clin Microbiol Infect Dis 11, 143–151, doi:10.1007/BF01967066 (1992).

25 Buck, C. B. et al. Carrageenan is a potent inhibitor of papillomavirus infection. PLoS Pathog 2, e69, doi:10.1371/journal.ppat.0020069 (2006).

26 Talarico, L. B. & Damonte, E. B. Interference in dengue virus adsorption and uncoating by carrageenans. Virology 363, 473–485, doi:10.1016/j.virol.2007.01.043 (2007).

27 Vives, R. R., Imberty, A., Sattentau, Q. J. & Lortat-Jacob, H. Heparan sulfate targets the HIV-1 envelope glycoprotein gp120 coreceptor binding site. J Biol Chem 280, 21353–21357, doi:10.1074/jbc.M500911200 (2005).

28 Moulard, M. et al. Selective interactions of polyanions with basic surfaces on human immunodeficiency virus type 1 gp120. J Virol 74, 1948–1960, doi:10.1128/jvi.74.4.1948-1960.2000 (2000).

29 Harrop, H. A. & Rider, C. C. Heparin and its derivatives bind to HIV-1 recombinant envelope glycoproteins, rather than to recombinant HIV-1 receptor, CD4. Glycobiology 8, 131–137, doi:10.1093/glycob/8.2.131 (1998).

30 Herold, B. C. et al. Poly(sodium 4-styrene sulfonate): an effective candidate topical antimicrobial for the prevention of sexually transmitted diseases. J Infect Dis 181, 770–773, doi:10.1086/315228 (2000).

31 Bourne, N., Zaneveld, L. J., Ward, J. A., Ireland, J. P. & Stanberry, L. R. Poly(sodium 4-styrene sulfonate): evaluation of a topical microbicide gel against herpes simplex virus type 2 and Chlamydia trachomatis infections in mice. Clin Microbiol Infect 9, 816–822, doi:10.1046/j.1469-0691.2003.00659.x (2003).

32 Van Damme, L. et al. Lack of effectiveness of cellulose sulfate gel for the prevention of vaginal HIV transmission. N Engl J Med 359, 463–472, doi:10.1056/NEJMoa0707957 (2008).

33 Leibbrandt, A. et al. Iota-carrageenan is a potent inhibitor of influenza A virus infection. PLoS One 5, e14320, doi:10.1371/journal.pone.0014320 (2010).

34 Eccles, R. et al. Efficacy and safety of an antiviral Iota-Carrageenan nasal spray: a randomized, double-blind, placebo-controlled exploratory study in volunteers with early symptoms of the common cold. Respir Res 11, 108, doi:10.1186/1465-9921-11-108 (2010).

35 Blakemore, W. R. Polysaccharide ingredients: carrageenan in Reference module in food science, doi:10.1016/B978-0-08-100596-5.03251-0 (Elsevier, 2015).

36 Sia, S. F. et al. Pathogenesis and transmission of SARS-CoV-2 in golden hamsters. Nature, doi:10.1038/s41586-020-2342-5 (2020).

37 Bao, L. et al. The pathogenicity of SARS-CoV-2 in hACE2 transgenic mice. Nature, doi:10.1038/s41586-020-2312-y (2020).

38 Li, H. et al. SARS-CoV-2 and viral sepsis: observations and hypotheses. Lancet 395, 1517–1520, doi:10.1016/S0140-6736(20)30920-X (2020).

39 Yang, Y. & Tang, H. Aberrant coagulation causes a hyper-inflammatory response in severe influenza pneumonia. Cell Mol Immunol 13, 432–442, doi:10.1038/cmi.2016.1 (2016).

