## Supplementary Figure S1 for "Antiviral activity of lambda-carrageenan against influenza viruses in mice and severe acute respiratory syndrome coronavirus 2 *in vitro*"

*** Corresponding author**

Meehyein Kim, Ph.D.

AdInfectious Diseases Therapeutic Research Center, Korea Research Institute of Chemical Technology, 141 Gajeongro, Yuseong, Daejeon 34114, Republic of Korea

[†] Current address: siRNAgen Therapeutics Co., Daejeon 34302, Republic of Korea

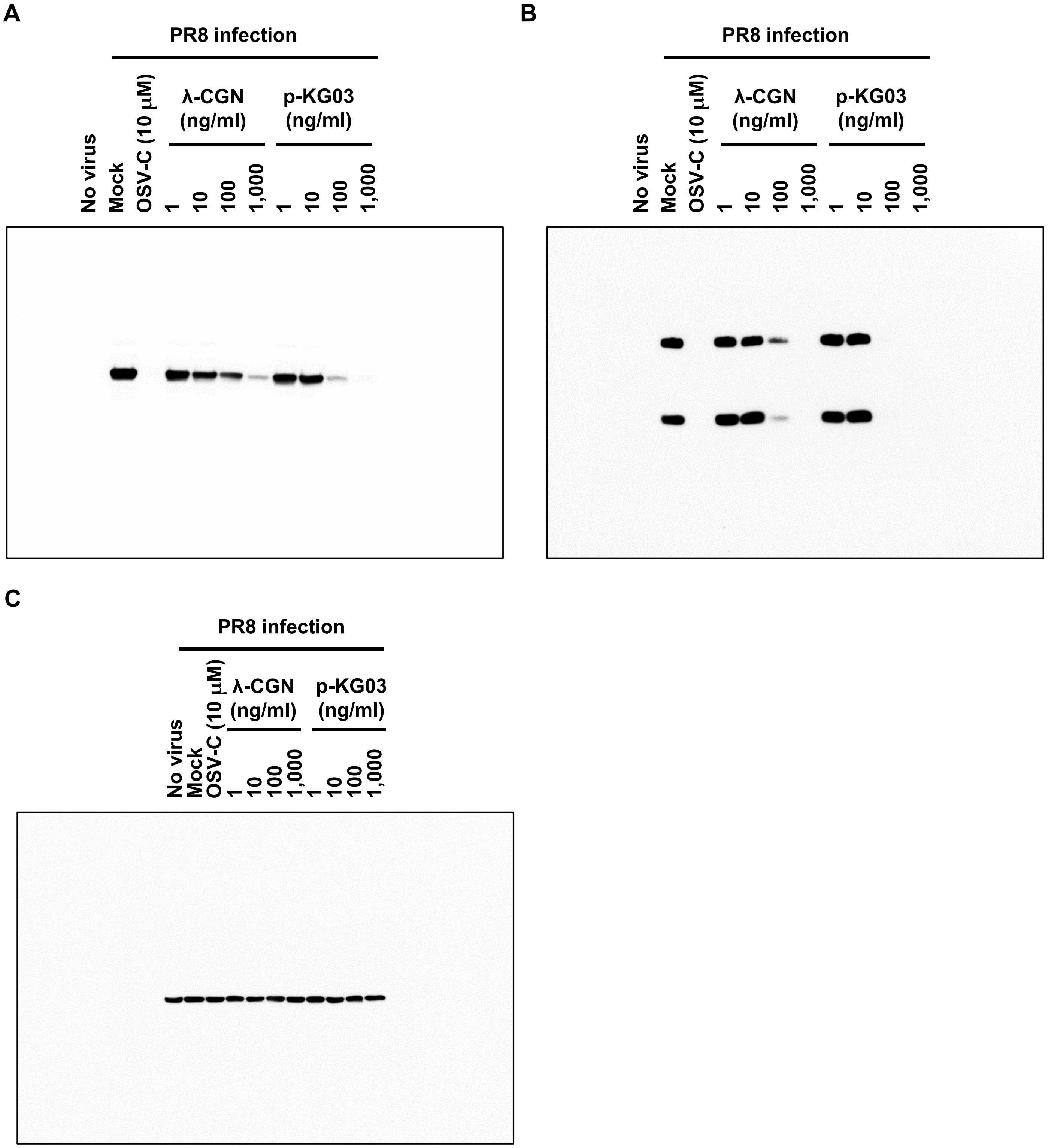


**Supplementary Figure S1.** Raw data showing full images of the western blots presented in Fig. 1B. MDCK cells infected with PR8 at an MOI of 0.001 were mock-treated (Mock) or treated with increasing concentrations of λ-CGN or p-KG03, or with 10 μM of OSV-C, at 35°C. On the next day, cell lysates were harvested for SDS-PAGE and immunoblotting with anti-NP (A) or anti-HA antibodies (B). β-Actin was used as a loading control (C). 'No virus' means negative control without viral infection. HRP on the membranes for detecting NP and HA was developed using Western Femto ECL kit (LPS Solution, Daejeon, Republic of Korea) (A, B), while it was done for β-actin using Super Signal West Pico Plus Chemiluminescent kit (Thermo Fisher Scientific, Rockford, IL, USA) (C).
